# Limits to adaptation in partially selfing species

**DOI:** 10.1101/026146

**Authors:** Matthew Hartfield, Sylvain Glémin

## Abstract

In outcrossing populations, “Haldane’s Sieve” states that recessive beneficial alleles are less likely to fix than dominant ones, because they are less expose to selection when rare. In contrast, selfing organisms are not subject to Haldane’s Sieve and are more likely to fix recessive types than outcrossers, as selfing rapidly creates homozygotes, increasing overall selection acting on mutations. However, longer homozygous tracts in selfers also reduces the ability of recombination to create new genotypes. It is unclear how these two effects influence overall adaptation rates in partially selfing organisms. Here, we calculate the fixation probability of beneficial alleles if there is an existing selective sweep in the population. We consider both the potential loss of the second beneficial mutation if it has a weaker advantage than the first, and the possible replacement of the initial allele if the second mutant is fitter. Overall, loss of weaker adaptive alleles during a first selective sweep has a larger impact on preventing fixation of both mutations in highly selfing organisms. Furthermore, the presence of linked mutations has two opposing effects on Haldane’s Sieve. First, recessive mutants are disproportionally likely to be lost in outcrossers, so it is likelier that dominant mutations will fix. Second, with elevated rates of adaptive mutation, selective interference annuls the advantage in selfing organisms of not suffering from Haldane’s Sieve; outcrossing organisms are more able to fix weak beneficial mutations of any dominance value. Overall, weakened recombination effects can greatly limit adaptation in selfing organisms.

## Introduction

Self-fertilisation – reproduction where both gametes arise from the same parent – frequently evolves from outcrossing species in nature. Self-fertilisation is widespread in angiosperms (Igic and Kohn 2006), some groups of animals (Jarne and Auld 2006) and fungi (Billiard *et al*. 2011; Gioti *et al*. 2012). It confers an initial benefit to an individual’s fecundity, including up to a 50% transmission advantage (Fisher 1941) and reproductive assurance under mate limitation (Baker 1955, 1967; Pannell *et al*. 2015). Both factors should allow selfing organisms to rapidly spread upon invasion of new habitats, unless countered by high levels of inbreeding depression (Lande and Schemske 1985). However, empirical studies usually find that selfing lineages are a ‘dead end’, since back-transitions to outcrossing are rare, and high extinction rates have been inferred from comparative studies of related selfing-outcrossing taxa (Igic *et al*. 2008; Goldberg *et al*. 2010; Wright and Barrett 2010; Wright *et al*. 2013).

Self-fertilisation has therefore been posited to be detrimental in the longterm. For an organism with selfing rate *σ*, the population has an inbreeding rate *F* = *σ*/(2 – *σ*), equivalent to Wright’s (1951) *F_IS_* statistic. The effective population size *N_e_* is reduced by a factor of at least 1/(1 + *F*) (Pollak 1987; Charlesworth 1992; Caballero and Hill 1992). Furthermore, the effective recombination rate is reduced in proportion to the inbreeding rate (Golding and Strobeck 1980; Nordborg 2000; Roze 2009). This joint reduction in both effective population size and recombination can lead to a decrease in the efficacy of selection. Deleterious mutations can therefore accumulate more rapidly in selfing organisms, leading to population extinction (Heller and Maynard Smith 1978; Lynch *et al*. 1995).

Whether this mechanism is a major cause of extinction of self-fertilising species is still under debate (reviewed in Glémin and Galtier (2012); Igic and Busch (2013); Hartfield (2016)). Some sister-species comparisons of selfing and outcrossing taxa reveal evidence of increased mutation accumulation in selfers, as demonstrated with either increased nonsynonymous-to-synonymous polymorphism ratio (*π_n_*/*π_s_*), or weaker codon usage bias. Conversely, analyses of divergence rates generally do not show evidence for relaxed selection. Part of the reason for this lack of evidence could arise due to recent transitions to selfing in most of these species, as explicitly demonstrated in *Capsella rubella* by Brandvain *et al*. (2013), thus leaving little time for mutation accumulation to act.

Less investigated is the idea that selfing reduces the ability of a species to adapt, especially to new environmental conditions, though it was the one initially formulated by Stebbins (1957). For adaptation at a single locus, selfing organisms are more likely than outcrossers to fix new recessive adaptive mutations (Haldane 1927; Charlesworth 1992) but are generally less efficient in adapting from standing variation (Glémin and Ronfort 2013). Yet the effect of adaptation at multiple loci in partially selfing organisms has received much less attention. Of particular interest is how the reduction in recombination efficacy in highly selfing organisms impedes the overall adaptation rate. A well-established phenomenon in low-recombining genomes is the ‘Hill-Robertson effect’, where the efficacy of selection acting on a focal mutation is reduced, due to simultaneous selection acting on linked loci (Hill and Robertson 1966; Charlesworth *et al*. 2009). Outcrossing can therefore break down these effects and unite beneficial mutations from different individuals into the same genome, greatly increasing the adaptation rate (Fisher 1930; Muller 1932; Felsenstein 1974; Otto and Barton 1997).

Historically, the effect of advantageous mutations on mating system evolution has been neglected, since most observable spontaneous mutations are deleterious in partial selfers (Slotte 2014), and the inbreeding depression they cause plays a central role in mating system evolution. Analyses using divergence data from the *Arabidopsis* genome shows low number of genes exhibiting signatures of positive selection (Barrier *et al*. 2003; Clark *et al*. 2007; Slotte *et al*. 2010, 2011), and only ~1% of genes have signatures of positive selection in *Medicago truncatula* (Paape *et al*. 2013). These analyses reflect broader findings that the proportion of adaptive substitutions in the coding regions of selfing plants is not significantly different from zero (Gossmann *et al*. 2010; Hough *et al*. 2013). However, widespread local adaptation to climate in *Arabidopsis* is observed (Fournier-Level *et al*. 2011; Hancock *et al*. 2011; Ågren *et al*. 2013), which is expected to leave a weaker signature on the genome at the species scale (Slotte 2014). However, the power to detect local selection can increase once demography and population structure are accounted for (Huber *et al*. 2014).

Finally, both outcrossing and selfing domesticated plant and crop species can also be used to demonstrate recent adaptation. Ronfort and Glémin (2013) showed how adaptive traits obtained from quantitative trait loci tended to be dominant in outcrossers and recessive in selfers, in line with ‘Haldane’s Sieve’ (where dominant mutants are more likely to fix than recessive types in outcrossers). Hence while beneficial mutations may not be as frequent as deleterious alleles, there exists evidence that they arise frequently enough to impact upon local adaptation and domestication in self-fertilising species. Furthermore, due to the reduced effective recombination rate in selfers, adaptive alleles should interfere with a greater region of the genome than in outcrossing organisms.

Recently, Hartfield and Glémin (2014) investigated the effect of a linked deleterious mutation on a selective sweep. In contrast to single-locus results, Hartfield and Glémin (2014) demonstrated how weakly-recessive beneficial alleles (i.e. those with *h* just less than 1/2) can contribute more to fitness increases in outcrossers than selfers, as they are less likely to fix deleterious alleles via hitchhiking. This model demonstrated how breaking apart selection interference at linked sites could provide an advantage to some degree of outcrossing, leading to mixed-mating being the optimal evolutionary state. A multi-locus simulation study by Kamran-Disfani and Agrawal (2014) verified that background selection impedes genome-wide adaptation rates in selfing organisms. These studies clearly showed how linkage to deleterious mutations can limit adaptation in selfers, yet it remains an open question as to what extent multiple beneficial mutations interfere in highly selfing species.

This article will extend previous analyses to consider how interference between beneficial mutations at linked sites affects their emergence in partially selfing species. Haploid two-locus analytical models of the Hill-Robertson effect are altered to take dominance and selfing into account, then examined to quantify how adaptation is affected.

## Outline of the problem

### General modelling approach

We wish to determine how the effect of existing beneficial mutations at linked loci impedes the fixation of novel adaptive alleles in partially selfing organisms. We consider two locus models to ensure tractability. Notation used in this study is outlined in Table 1.

**Table 1:** Glossary of Notation.

| Symbol | Usage |
| --- | --- |
| $2N$ | Number of haplotypes in the population (of size $N$) |
| $A, B$ | Locus at which first, second beneficial allele arises |
| $r$ | Recombination rate between two loci |
| $s_A, s_B$ | Fitness coefficients of original and new advantageous alleles |
| $h_A, h_B$ | Dominance coefficients of original and new advantageous alleles |
| $p$ | Frequency of first advantageous allele at timepoint |
| $q$ | Frequency of second advantageous allele at timepoint (if $s_A < s_B$) |
| $\sigma$ | Proportion of matings that are self-fertilising |
| $F$ | WRIGHT'S (1951) inbreeding coefficient, $\sigma/(2 - \sigma)$ |
| $\Phi$ | Probability of identity by descent at two loci (ROZE 2009) |
| $P_A(p), P_B(p)$ | Fixation probability of original and new allele if unaffected by linkage (Equation 1) |
| $P_{AB}^*$ | Fixation probability of both mutants in absence of interference |
| $P_{AB}(p)$ | Actual fixation probability of both mutants, after accounting for interference |
| $\bar{P}_{AB}$ | Average $P_{AB}$ over one or several selected substitutions at locus $A$ |
| $R(p)$ | Ratio of actual to non-interference double-allele fixation probability, $P_{AB}/P_{AB}^*$ |
| $\bar{R}$ | Average $R$ over one or several selected substitutions at locus $A$ |
| $P_{HH}$ | Fixation probability of second allele with wildtype background |
| $P_d(q)$ | Fixation probability of haplotype carrying both sweeps |
| $\tau$ | Time taken for first sweep to reach frequency $p$ |
| $T$ | Scaled time, $s_A t$ |
| $\Pi(p)$ | Average fixation probability of second allele if it does not replace the first sweep |
| $\Pi_{rep}(p)$ | Probability that second sweep replaces first if $s_B > s_A$ |
| $Q_0(p), Q_1(p)$ | Fixation probability of novel allele if appearing on neutral or already beneficial genetic background |
| $\theta_1(p), \theta_0(p)$ | Relative selective advantage of second allele, if residing on either beneficial or neutral background |
| $\theta_d(q)$ | Relative selective advantage of recombinant haplotype carrying both alleles |
| $\Delta$ | Difference in fixation probability between different backgrounds, $Q_1 - Q_0$ |
| $\phi$ | Scaled advantage of new beneficial allele, $s_B/s_A$ |
| $\rho$ | Scaled recombination rate, $r/s_A$ |
| $\lambda_A$ | Rate of selected substitution at locus $A$ |
| $\Theta$ | Population rate of beneficial mutation at single locus, $4Nu_a$ |

We assume a diploid population of fixed size *N*, so there are 2*N* haplotypes present. Each haplotype consists of two liked loci with recombination occurring at a rate *r* between them. At a first locus *A*, consider a beneficial mutation *A*_1_ with selective coefficient *s_A_*, so the fitness of individuals carrying it is 1 + *h_A_s_A_* in heterozygote form, and 1 + *s_A_* in homozygote form. Similarly at a linked locus *B*, the fitness of individuals carrying the beneficial mutation *B*_*1*_ is 1 + *h_B_s_B_* in heterozygotes and 1 + *s_B_* in homozygotes. The wild-type alleles are denoted *A*_0_, *B*_0_; the four haplotypes are *A*_0_*B*_0_, *A*_1_*B*_0_, *A*_0_*B*_1_ and *A*_1_*B*_1_. We further assume that selection acts on genotypes at the diploid stage and that fitness between loci is additive. As an example, an individual composed of the genotype *A*_1_*B*_0_/*A*_1_*B*_1_ will have fitness 1 + *s_A_* + *h_B_s_B_*.

For strongly selected mutants (assuming 2*N_e_hs* ≫ 1), the trajectory of a beneficial mutation can be decomposed into three parts. A schematic is shown in Figure 1. The stages are (i) a initial stochastic phase at low frequency where extinction by drift is likely; (ii) conditioned on escaping initial extinction (i.e. emergence), the allele increases in frequency on a quasi-deterministic trajectory; (iii) a second stochastic phase at very high frequency where fixation is almost certain (Kaplan *et al*. 1989). If two mutations segregate simultaneously at low frequency in the stochastic zone, they do not influence each other and their fates can be assumed to be independent. However, as soon as one mutant has emerged and started to sweep quasi-deterministically it affects the fate of the other mutation. When considering a single unlinked mutation, once it has emerged its ultimate fixation is almost certain (which corresponds to the branching process approximation). The probability of fixation is thus equal to the probability of emergence. However, when two (or more) mutations interfere, a mutation that has emerged can be replaced by a linked competitor and ultimately lost, which is well known in asexual species as the ‘clonal interference’ effect (Gerrish and Lenski 1998). If so, the probability of fixation can be lower than the probability of emergence. Under tight linkage, or with a high selfing rate (as the loss of heterozygosity will reduce the effective recombination rate), this process has to be taken into account.

**Figure 1:**
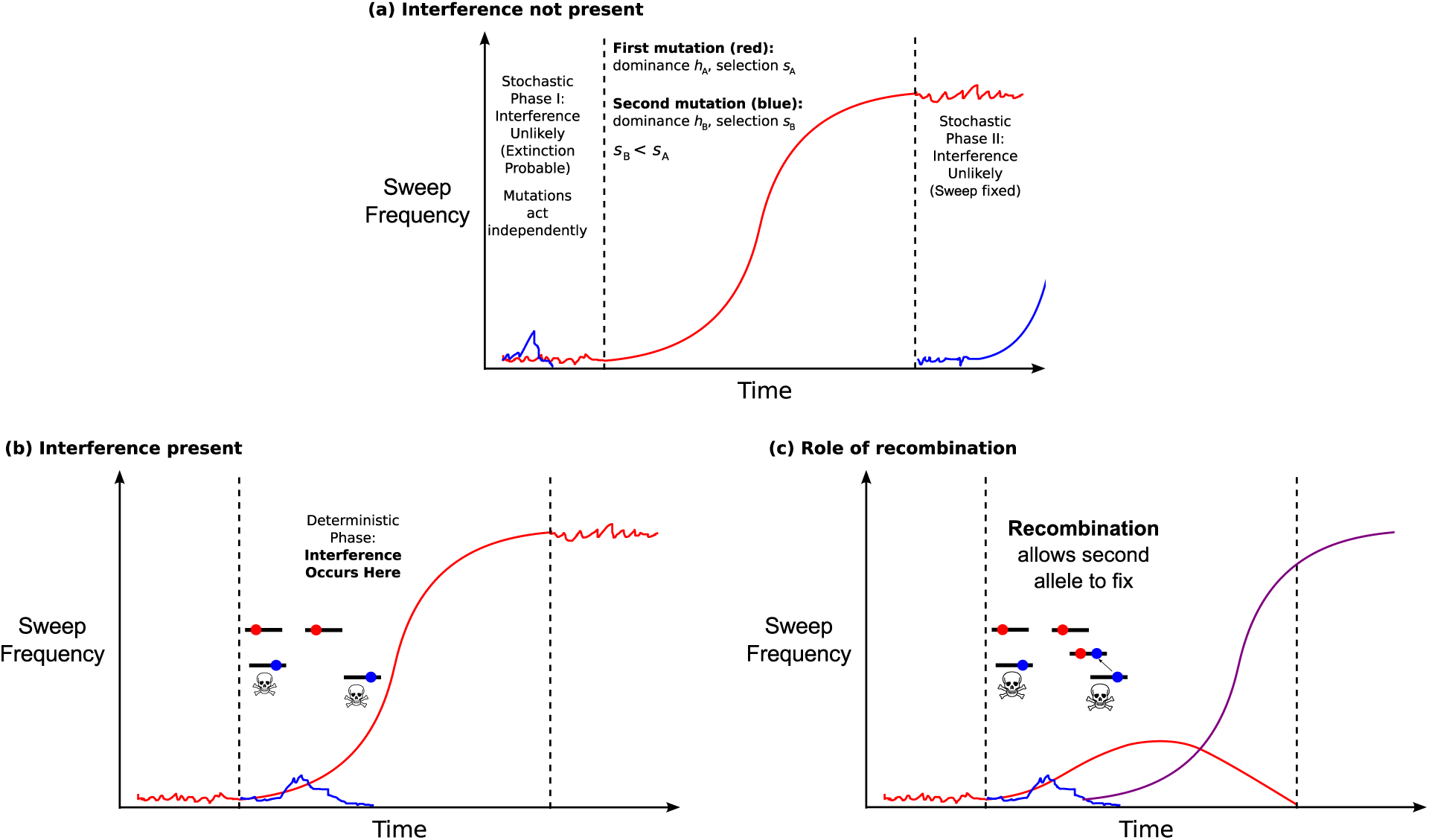
Schematic of the model for the case *s_B_* < *s_A_*, i.e. the second mutant is weaker than the first. We assume *h_A_* = *h_B_* here for simplicity. (a) The red line denotes the frequency of the initial beneficial mutation over time. When it is at low (*p* ~ 1/2*N*) and high (*p* ~ (1 – 1/2*N*)) frequencies, changes in frequency are determined stochastically. Any linked beneficial alleles that appear during these phases will act independently as one genetic background dominates that the second mutant can appear on. (b) Once the first allele is sufficiently prevalent, it will increase in frequency over time in a regular manner (the ‘deterministic’ phase). Any secondary alleles that appear during this phase will be impacted by selection interference, so will be less likely to fix even if they do emerge. These alleles are represented by blue dots, with the trajectory shown by a blue line. (c) Interference can be broken down if recombination moves the blue allele onto the fitter background containing the red allele, and this new haplotype (whose frequency is shown by a purple line) emerges in the population.

We assume that mutation *A*_1_ is the first to reach a high copy number and escape extinction by drift (although it could have been the second to arise), so is segregating in the population. Its trajectory can be modelled using deterministic equations. Without interference, the probability of fixation of the two mutations, 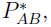, is simply equal to the single-locus probability of fixation of the second mutation:

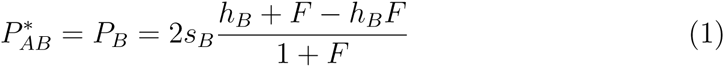

(Caballero and Hill 1992; Charlesworth 1992). 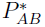 can also be defined as the fixation of both alleles given that *A*_1_ has already fixed in the population, hence *P_A_* does not appear. Equation 1 leads to the classical result that the probability of fixation is higher under outcrossing than under selfing when *h_B_* > 1/2. More generally, the emergence of mutation *B*_1_ depends on the genetic background it appears on, and the switching rate between backgrounds through recombination, which is the cause of the ‘Hill-Robertson’ effect we wish to model (Hill and Robertson 1966). Denoting the actual fixation probability of both mutants as *P_AB_*(*p*), we then define the degree of interference *R*(*p*) as the ratio 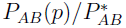. measures to what extent fixation probability is reduced due to the presence of linked mutation. For example, *R*(*p*) = 0.1 means that the fixation probability of a linked mutant is a tenth that it would be if unlinked, given the first allele is at frequency *p*. Hence it is a measure of how severe selection interference is at impeding fixation of subsequent beneficial mutations.

In the simplifying case *h_A_* = *h_B_*, the dynamics of how the second mutation emerges will differ depending on whether *s_B_* < *s_A_* or vice versa. Hence a key parameter in the full model is the ratio *ϕ* = *s_B_*/*s_A_*. If *s_B_* < *s_A_* (*ϕ* < 1), the dynamics of the first mutation is not influenced by the second and cannot be replaced. We thus only need to compute the emergence probability of the second mutation, which is likely to go extinct unless it appears on or recombines onto the background carrying the first beneficial mutation. Barton (1995) outlined a general model to calculate this effect for a haploid organism. We will demonstrate how diploidy and selfing can be accounted for in that model and subsequently compute Π(*p*), which is the contribution to R(*p*) arising from selection interference alone.

If *s_A_* < *s_B_* (*ϕ* > 1) then the second mutation can replace the first if it arises on the wild-type background, and no successful recombinant occurs. We can calculate the probability of this effect by adjusting the analysis of Hartfield and Glémin (2014) to consider two beneficial mutations. We thus need to subtract from Π(*p*) the probability that mutation *B*_1_ replaces mutation *A*_1_ once it has emerged, denoted by Π_*rep*_(*p*). In the general case, the degree of interference will be given by:

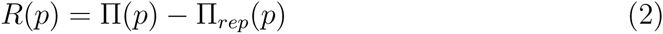

In practice, instead of calculating *R*(*p*), we instead measure *R̅*. This is the *R* quantity for mutation *B*_1_ arising when mutation *A*_1_ is at a given frequency *p*, averaged over the whole possible origin times of the second mutation. Formal definitions of these conditions are presented below.

### A simple first analysis: complete selfing versus outcrossing with free recombination

Before deriving the full model, we can compare the two most extreme cases that can be investigated. Under outcrossing and free recombination, the fates of the two mutations are essentially independent. Hence the fixation probability of the second mutation, conditioned on the first having emerged, is simply the single locus probability of fixation given by Equation 1 with *F* = 0 (Haldane 1927). At the other extreme with complete selfing (*F* = 1), recombination is suppressed so interference is maximised. Here, a second mutation can only fix if it appears on the same genetic background as the original beneficial allele, which is present at frequency *p*. Previous theory (Hartfield and Otto 2011; Hartfield and Glémin 2014) on emergence in this scenario gives the fixation probability of the double mutant as:

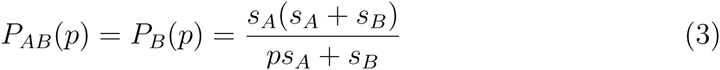

See Equation 7 of Hartfield and Glémin (2014) with *s_d_* = *s_A_* and *s_A_* = *s_A_* + *s_B_*. The mean probability of fixation of both alleles thus involves integrating Equation 3 over the entire sweep, assuming that the second mutation arises at a time that is uniformly distributed during the first sweep:

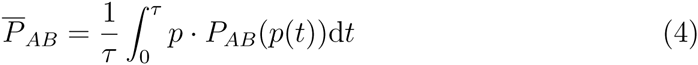

where *τ* is the duration of the first sweep. We can also solve Equation 4 over *p* from *p*_0_ to 1 – *p*_0_; the term inside the integral is divided by *d_p_*/*d_t_* = *s_Ap_*(1 – *p*) to remove time dependence. Solving in the limit of large population size (i.e. *p*_0_ = 1/(2*Ns*) → 0) leads to *P̅_AB_* = *s_B_*/2 (see Supplementary Mathematica File S2): full linkage reduces the emergence probability by a half (*R̅* = 1/2). Intuitively, this can be explained by the fact that as population size increases, the deterministic phase of the first sweep becomes shorter compared to the initial and final stochastic phases 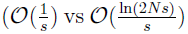; Ewing *et al*. (2011)). The second mutation thus occurs roughly half of the time during the initial stochastic phase where its probability of arising on the beneficial background, hence of emerging, is very low (*p* ≈ 0 in Equation 4). Alternatively, it can appear half of the time during the last stochastic phase where it almost always originates in the beneficial background and its probability of emerging is approximately *s_B_* (*p* ≈ 1 in Equation 4).

By comparing this result to that with outcrossing and free recombination (2*h_B_s_B_*), outcrossing is more able to fix both mutants if *h_B_* > 1/4, instead of *h_B_* > 1/2 without interference. However, the advantage to outcrossing may not be as high, since the true degree of inference depends on the strength of both mutations and the recombination rate. In addition, the degree of stochastic interference also depends on the flow of beneficial mutations, which depends on the mating system. We now turn to the full model to exactly quantify this effect.

## Modelling Framework

### Deriving the reduction in emergence probability due to interference

We first need to determine Π(*p*), the reduction in the emergence probability of the second mutation when it arises, given the first is at frequency *p*. We use branching process methods for calculating mutation emergence if acting over multiple genetic backgrounds. In a seminal paper, Barton (1995) outlined how to calculate the emergence probability of a focal beneficial allele that changes between different backgrounds in a haploid population. If the probability of switching between backgrounds is of the same order as selection coefficients, *s*, and difference in emergence probability over backgrounds is of order *s*^2^, Barton (1995) showed that the emergence probability of a novel beneficial allele in background *i* at time *t, Q_i_*, is a solution to the following differential equation:

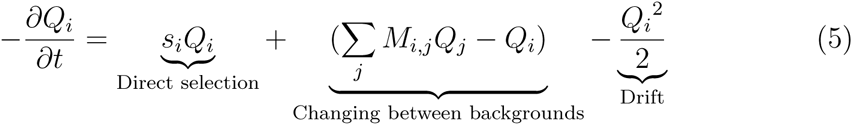

The right-hand side of Equation 5 can be decomposed into three terms. The first is a direct selection term: the fixation probability increases if the beneficial allele has a higher fitness advantage in background *i, s_i_*. *M*_*i,j*_ is the probability that offspring in background *i* moves to background *j* per generation; in this case, this effect arises from recombination changing the genetic background of the focal allele. Finally, there is a negative 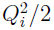 term, denoting how genetic drift can cause the allele to go extinct. Note that the differential equation term is negative, since this system of equations is considered going back in time. Barton (1995) subsequently used this framework to investigate the fixation probability of a second beneficial allele, given that it arises in close linkage to an existing sweep.

Our goal is to derive equivalent equations for a diploid, self-fertilising population. In Supplementary File S1 and the Supplementary Mathematica File S2, we show how similar equations can be derived for outcrossing species. However, when selfing is including then the full system of equations becomes unwieldy. In order to obtain analytical solutions, we proceed by using a separation-of-timescale argument. When recombination is low, as assumed in Equation 5, haplotypes carrying either the first beneficial allele or the wildtype quickly reach their genotypic equilibria with inbreeding (Wright 1951); specifically, we assume that equilibrium is reached between recombination events. If selection acts on genotypes, we can use the relative fitness advantage of each genotype to determine that of the second beneficial allele, depending on the haplotype it appears on. We can then create a variant of Equation 5 with these steady-state values. Using this argument, we obtain a tractable form for the transition probabilities between backgrounds (Supplementary File S1).

With high selfing rates, Equation 5 remains valid for any recombination rate as the probability of moving from one background to another can be low. But as sweep effects can span large genetic map distances under high selfing, it is important to analyse this special case properly. Roze (2009, 2015) derived the equilibrium genotype frequencies at two loci under partial selfing as a function of the probability of identity by descent at a single locus, *F*, and at two loci, Φ. This approach takes into account correlations of homozygosity between linked loci. In Supplementary File S1 and Mathematica File S2, we show that Φ equilibrates as quickly as *F*, so the separation-of-timescale argument can be used for any recombination level, assuming that two-locus equilibrium genotype frequencies for given selfing and recombination rates are instantaneously reached, compared to change in allele frequencies. This approximation should work well at equilibrium, but not necessary in non-equilibrium conditions such as during selective sweeps. However, because equilibrium values are quickly reached, these calculations are also accurate under more general conditions (see simulation results below).

To complete the model, we re-derive each component of Equation 5 in turn to account for diploidy and selfing.

**Direct selection of the second allele.** Let *Q*_1_(*p*) denote the probability that the new allele fixes, given that it arises in linkage with the existing mutant *A*_1_ (which is at frequency *p*). *Q*_0_(*p*) is the fixation probability if the second allele appears on the wild-type (neutral) background. We further denote the relative selective advantage of each haplotype (either containing both advantageous alleles, or the second allele only) by *θ*_1_(*p*) and *θ*_0_(*p*). These are derived by calculating the frequency of each potential haplotype background that the second allele appears on, and the relative proportions of each. Calculations are outlined in Supplementary File S1:

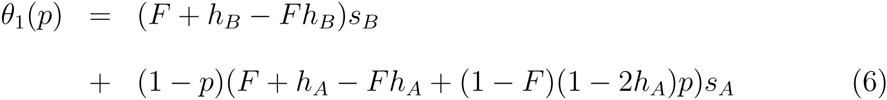

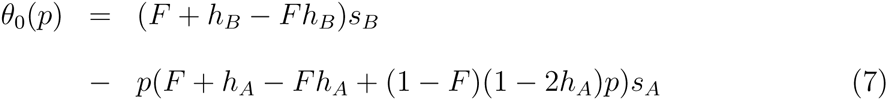

Note that Equations 6 and 7 are the same for both the general and the low recombination cases. As in Barton (1995) the trajectory of the first mutation *A*_1_ is assumed to be deterministic, and hence described by the following differential equation:

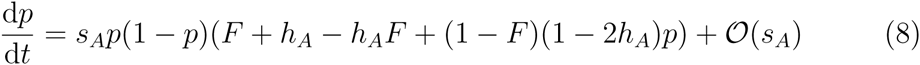

Furthermore we can rescale time by selection, i.e. setting *T* = *s_A_t* (Barton 1995).

**Rescaling recombination.** Nordborg (2000) used a coalescent argument to show that in selfing populations, when recombination is of order 1/2N, the recombination rate is scaled by a factor (1 – *F*) (see also Golding and Strobeck (1980)). This arises as a proportion *F* of recombination events are instantly ‘repaired’ due to selfing. We show in Supplementary Mathematica File S2 that the same result can be obtained by considering the decay of linkage disequilibrium in an infinite population with the less restrictive condition of *r* being small (*r* ≪ 1) but not necessarily ***O***(1/2*N*). However, this scaling breaks down for high recombination and selfing rates, as it becomes likely that recombination occurs between genotypes within individual lineages (Padhukasahasram *et al*. 2008). Relaxing the assumption of low recombination, we can derive a more exact rescaling term following Roze (2009, 2015): *r*(1 – 2*F* + Φ), which reduces to the coalescent rescaling *r*(1 – *F*) with low recombination. The equilibrium value for Φ can then be used (see Equation A8 in Supplementary File S1, and Roze and Lenormand (2005); Roze (2009)). Furthermore, in Supplementary Mathematica File S2 we show that the error will be at most a factor 2/(2 + *F*) with high recombination. Given that the effective recombination rate will remain small with high selfing rates irrespective of the scaling term, using the coalescent rescaling should capture the average fixation probability, but more accurate expression can be obtained using *r*(1 – 2*F* + Φ) instead.

**Creating the system of equations.** Hence the system of equations in Barton (1995) are modified to:

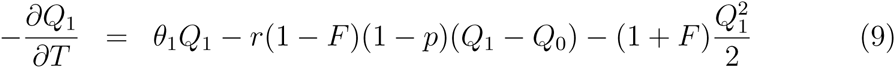

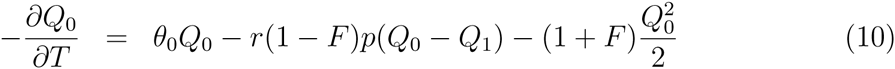

Equations 9 and 10 account for the relative selective advantage of the second allele, *θ* (with *p* verifying Equation 8); the decrease in effective recombination rate *r*(1 – *F*) (or *r*(1 – 2F + Φ) for greater accuracy); and the *Q*^2^/2 terms are scaled by 1 + *F* due to an increase in genetic drift (Pollak 1987; Charlesworth 1992; Caballero and Hill 1992).

Calculating Π. We next follow the approach of Barton (1995) and investigate the average fixation probability over haplotypes given the first beneficial allele is at a certain frequency, defined as Π = *p*^*Q*^_1_ + (1 – *p*)*Q*_0_, and the difference in emergence probability between the backgrounds, Δ = *Q*_1_ – *Q*_0_. These terms are scaled by the probability of fixation of the second allele if unlinked, (2*s_B_*(*F* + *h_B_* — *Fh_B_*))/(1 + *F*), so Π lies between 0 and 1. We also introduce the rescaled parameters *ϕ* = *s_B_*/*s_A_* and *ρ* = *r*/*s_A_* to determine how the relative selective strengths and recombination rates affect interference. We thus obtain:

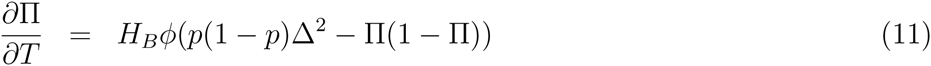

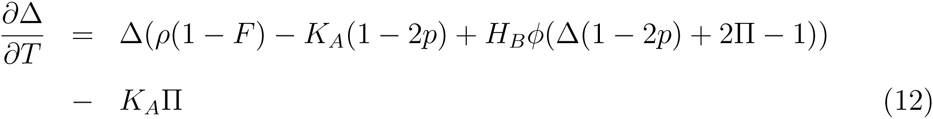

where *H_B_* = *h_B_* + *F* – *h_B_F* and *K_A_* = *h_A_* + *F* – *h_A_* + (1 – *F*)(1 – 2*h_A_*)*p*. Note that if we use the more exact recombination rescaling term *r*(1 – 2*F* + Φ), it is not possible to fully factor out *s_A_* from Equations 9 and 10 as Φ is a function of *r*.

For a given time of origin of the second mutation, *t*, the joint solution of this system and Equation 8, 11 and 12 gives Π(*p*(*t*)). These equations must be solved numerically by, e.g., using the ‘NDSolve’ function in *Mathematica* (Wolfram Research, Inc. 2014). Alternatively, to remove the time dependence (*dt)* and directly obtain Π(*p*), we can divide both Equations 11 and 12 by *d_p_*/*d_t_* (Equation 8). Boundary conditions can be found by looking at the behaviour of the system as *t* → ∞ or *p* → 1. In this case, we observe that Π → 1, reflective of the fact that as the first mutation fixes, the second allele is certain to arise alongside it. Hence the second allele’s fixation probability is not reduced. Boundary conditions for Δ can be calculated by assuming *ϕ* ≪ 1 (as used in Barton (1995)) and *∂*Δ/*∂T*/ → 0 as *p* → 1. In this case Δ tends to (1 – (1 – *F*)*h_A_*)/(1 – (1 – *F*)(*h_A_* – ρ)), which reflects the probability that the second allele can recombine onto the fitter background if appearing on a wild-type chromosome, otherwise it is guaranteed to be lost (Barton 1995). Although this condition assumes small *ϕ*, the system of equations appear to work well even with larger *ϕ* when compared to simulations.

### Integration over the sweep trajectory

To obtain the average effect of interference we need to consider all possible origins of the second mutation. The average *R* for mutation *B*_1_ arising uniformly in a long time interval [*T*_0_,*T*_1_] spanning the sojourn of mutation *A*_1_ is given by:

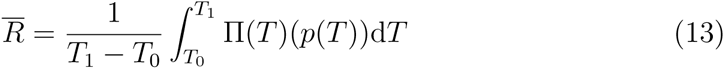

As previously showed by Barton (1995), *R̅* can be approximated by:

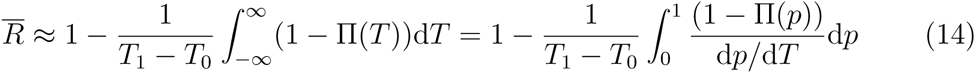

Integration from very ancient time (equivalently, a frequency lower than 1/2N) reflects the fact that mutation *A*_1_ affects the fate of mutation *B*_1_ even if it appears afterwards, when mutation *B*_1_ is still at a low frequency in the stochastic zone. Hence it is not obvious if a natural choice for *T*_1_ – *T*_0_ is the sojourn time of the first mutation. Moreover, because selfing and dominance affect the fixation time of alleles (Glémin 2012), averaging over this time would not allow direct comparison between different selfing rates and dominance levels. For example, the effect of linked mutation is expected to be stronger under selfing than under outcrossing but the time interval when interference can occur is shorter. Finally, interference also depends on the substitution rate at locus *A*, which is also affected by selfing and dominance. All these effects can be taken into account by assuming a steady state of substitutions at a low rate at locus *A* (i.e. no multiple substitutions). The rate of selected substitution at locus *A* is given by:

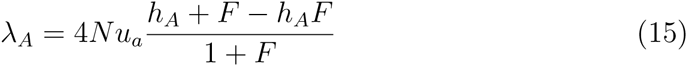

where time is measured in 1/*s_A_* generations. Following Barton (1995) we use:

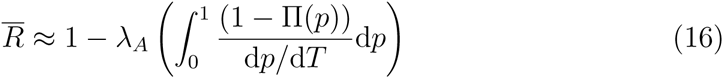

The justification is as follows. The waiting time between two sweeps is exponentially distributed with mean 1/*λ_A_*. If *T*_1_ – *T*_0_ < 1/*λ_A_*, interference between mutation *A*_1_ and *B*_1_ thus occurs for a proportion of time (*T*_l_ – *T*_0_)/(1/*λ_A_*). On average, the effect of *A*_1_ on *B*_1_ is thus:

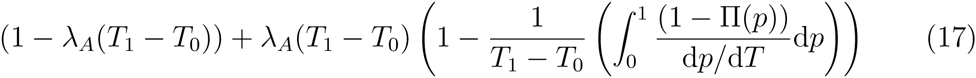

leading to Equation 16.

### Deriving the probability of sweep replacement, Π_*rep*_

The previous analysis focussed primarily on the case where the second mutant is weaker than the first, so *R*(*p*) = Π(*p*). However, if selection acting on *B*_1_ is sufficiently strong then it is possible that *B*_1_ replaces *A*_1_, so only *B*_1_ fixes. We need to calculate the probability of this replacement occurring and subtract it from the baseline reduction Π(*p*). This probability can be calculated by altering the model of Hartfield and Glémin (2014), which investigated a deleterious allele hitchhiking with a sweep. In our case, the ‘deleterious’ allele is the wildtype allele at the first locus *A*_0_, and the ‘advantageous’ allele the second fitter sweep *B*_1_. Hartfield and Otto (2011) implemented a similar rescaling for a haploid model, while Yu and Etheridge (2010) provided a general stochastic algorithm for investigating this behaviour.

A fuller derivation is included in Supplementary File S1. By calculating *R*(*p*) from first principles, we can infer that:

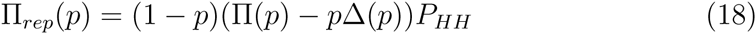
where Π(*p*) and Δ(*p*) are given by Equations 11 and 12. Equation 6 of Hartfield and Glémin (2014) is then used to calculate *P_HH_*:

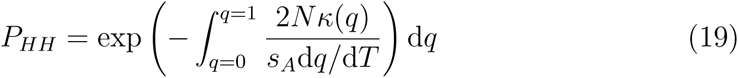

for *k*(*q*) = *q*(1 – *q*)*r*(1 – *F*)*P_d_*(*q*) where *q* is the frequency of allele *B*_1_. As above, *r*(1 – 2*F* + Φ) can be used instead of *r*(1 – *F*) to create more precise expressions. *P_d_*(*q*) is the emergence probability of the haplotype carrying both beneficial alleles (*A*_1_*B*_1_) if it formed by recombination. It is the solution of the following equation, where *θ_d_* is the relative fitness of this haplotype:

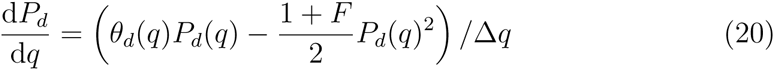

The mean interference effect is similar to Equation 16. However, contrary to emergence, the replacement of mutation *A*_1_ by mutation *B*_1_ can occur only if mutation *B*_1_ arises when mutation *A*_1_ has already emerged; that is, for *p* > *p_e_* ≈ (1 + *F*)/[2*N_s__A_*(*h_A_* + *F* – *h_A_F*)]. This condition is a bit too restrictive because we should also consider the case when mutation *A*_1_ arises after but emerges before mutation *B*_1_. Moreover, the distribution of *p_e_* should be used instead of the average value. However, these complications have only minor quantitative effects (not shown). If considering replacement, Equation 16 is written as:

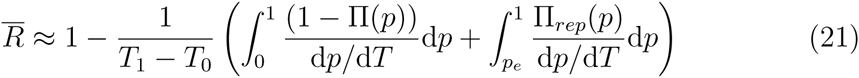

## Simulations

We tested the accuracy of analytical solutions by comparing them to stochastic simulations written in C; code is available as Supplementary File S3 and online from http://github.com/MattHartfield/TwoAdvSelfSims. When measuring Π, the first allele was seeded at initial frequency *p*; given this frequency and selfing rate *σ*, the proportion of mutant homozygotes, heterozygotes, and wild-type homozygotes were calculated based on standard equations with inbreeding (Wright 1951). The second allele was subsequently introduced onto a random background with frequency 1/2*N* (i.e. as a single copy). Frequencies of each genotype were altered deterministically by a factor *w_g_*/*w̅* due to selection, where *w_g_* is the fitness of each genotype and *w* is the population mean fitness. Recursion equations derived by Hedrick (1980, Equation 3) then calculated how genotype frequencies changed due to partial selfing. A life-cycle was completed by resampling *N* genotypes from a multinomial distribution to implement random drift. The second allele was tracked until one haplotype fixed, with the simulation repeated until 5000 fixations of the second beneficial allele occurred. It was noted how often each haplotype fixed; from this data we subsequently calculated the second allele fixation probability, relative to the expected result without interference. When measuring *P_HH_* we instead measured how often the haplotype carrying solely the fitter mutant fixed. Confidence intervals were calculated using the method of Agresti and Coull (1998).

## Results

### Validating of the analytical approach

**Testing Π**. We first tested the accuracy of Π, as given by Equation 11, with stochastic simulations. A subset of comparisons are shown in Figure 2; fuller comparisons are given in Supplementary Mathematica File S2. We see that on the whole, the analytical solutions provide an accurate match with simulations for a wide variety of selfing and dominance values. This includes cases of high selfing and recombination, despite our model assuming low recombination. Nevertheless, simulation results mildly underestimate the original analytical solutions with very high values (e.g. *F* = 0.99, *r* > 0.2; see Figure 2(*f*)). Using the more exact rescaled recombination rate, *r*(1 – 2*F* + Φ), improves the fit. Some inaccuracies are also apparent if the first allele is at a low frequency when the second appears (p ≈ 0.01). This discrepancy likely arises due to the trajectory of the first beneficial allele not being completely deterministic when starting at low frequency.

**Figure 2:**
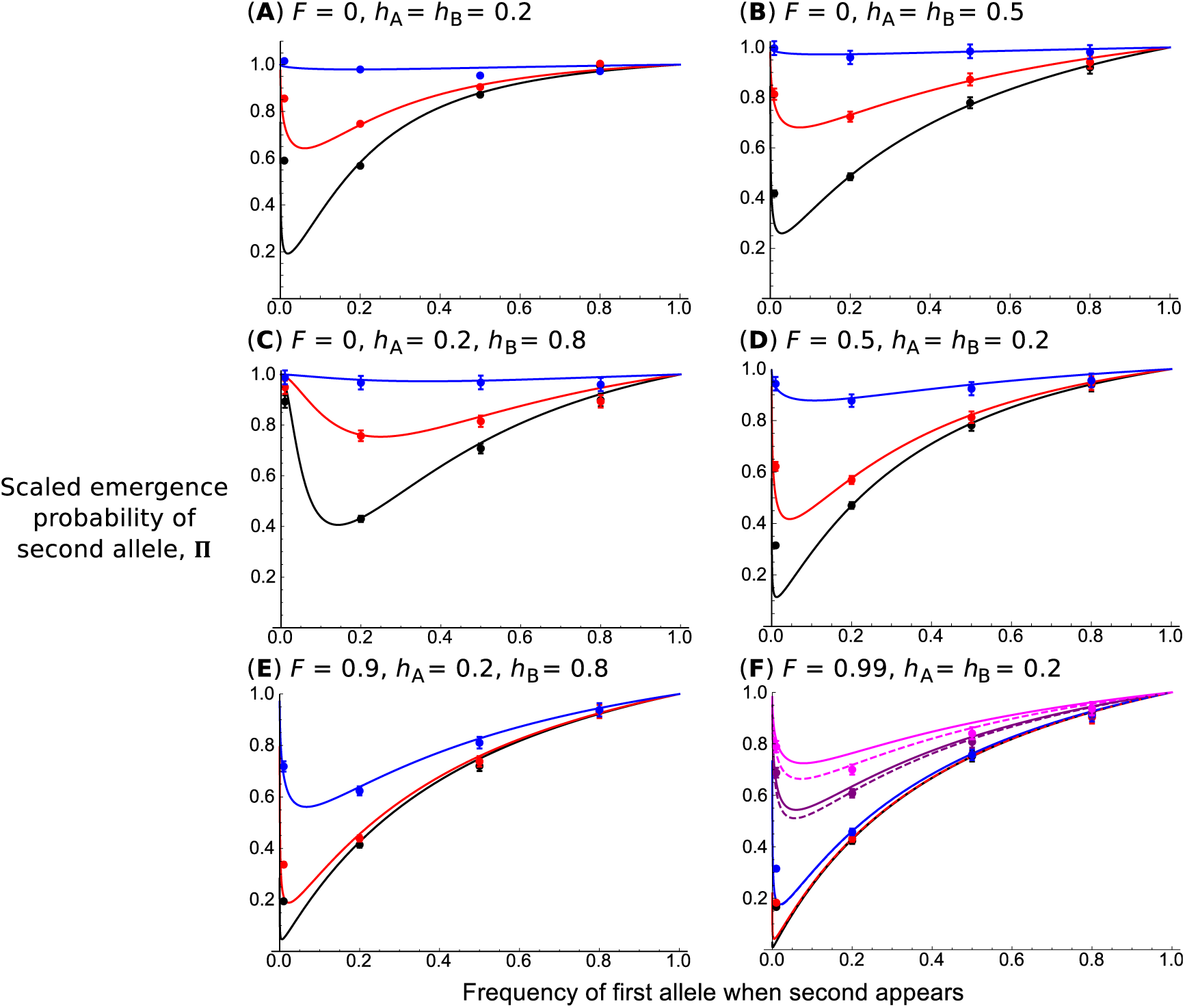
Probability of fixation of the second allele relative to the unlinked case, Π, as a function of the first allele frequency, *p*. *N* = 10, 000, *s_A_* = 0.01, *s_A_* = 0.005 (so *ϕ* = 0.5), and from bottom to top in (a)-(d): *r* = 0.0001, 0.001, 0.01 (corresponding to *ρ* = 0.01, 0.1, 1). In (f), there are also *r* = 0.1 and 0.25 results added (corresponding to *ρ* = 10 and 25). Other parameters used are listed above each subplot. Curves correspond to solutions provided by analytical system of differential equations (Equation 11), rescaled so it is a function of *p* instead. In (f), dashed lines are analytical results using the more exact recombination rate, *r*(1 – 2*F* + Φ). Points corresponds to 5,000 stochastic simulations for which the second beneficial allele has fixed. Bars represent 95% confidence intervals; if they intervals cannot be seen, they lie within the plotted points. Note that in panel (a), simulation results are presented after rescaling fixation probability by the diffusion equation solution, to account for recessive alleles (see main text for details).

In addition, if the second allele is highly recessive where there is outcrossing (*h_B_* = 0.2), the simulated scaled allele fixation probability can be higher than in single-loci models. This is simply because the fixation probability of recessive beneficial mutants are underestimated using the branching-process solution without considering homozygote genotypes (Equation 1, which holds for highly recessive alleles only in very large population sizes, i.e. at least *N* = 100, 000). For smaller population sizes a diffusion-equation solution, *P_dif_*, offers the correct baseline emergence probability (Caballero and Hill 1992). Hence rescaling the *h_B_* = 0.2 simulations by this solution, *P_dif_* Π (instead of *P_B_*Π where *P_B_* is given by Equation 1) causes simulations to match with analytical solutions.

**Testing Π_*rep*_.** Figure 3 shows the estimate of Π_*rep*_ compared to simulation data if *s_B_* > *s_A_* and the first sweep is at frequency p. Generally, if the first mutation is not recessive, recombination is low (and/or selfing high) and *p* not too low (greater than 1/*N*) then the analytical solution matches up well with simulations. The fit is not improved using the *r*(1 – 2*F* + Φ) recombination term, since the unscaled recombination rate remains low (*r* ≪ 1). However, if recombination increases (2*Nr* approaches 1) and mutations are recessive, then the actual replacement probability can be underestimated (for example, with *h_A_* = *h_B_* = 0.2; Figure 3(b)). By tracking the frequencies of individual haplotypes over time, we can determine that in cases where the model fails, it is because two key assumptions are violated (Supplementary Mathematica File S2). In particular, we assumed that recombination only occurs between haplotypes carrying one of the beneficial alleles. But in this case, the wild-type haplotype is not rapidly eliminated. Hence only a fraction of recombination events occurs between haplotypes carrying beneficial alleles, and Equation 19 would overestimate the effect of recombination. This error would not be large if net recombination is low. Furthermore, the first beneficial allele does not increase in frequency at the start of the process. This behaviour violates the assumption that it will compete with the second allele. These modelling violations are also observed if both alleles are dominant in outcrossing populations (*h_A_* = *h_B_* = 0.8; see Supplementary Mathematica File S2).

**Figure 3:**
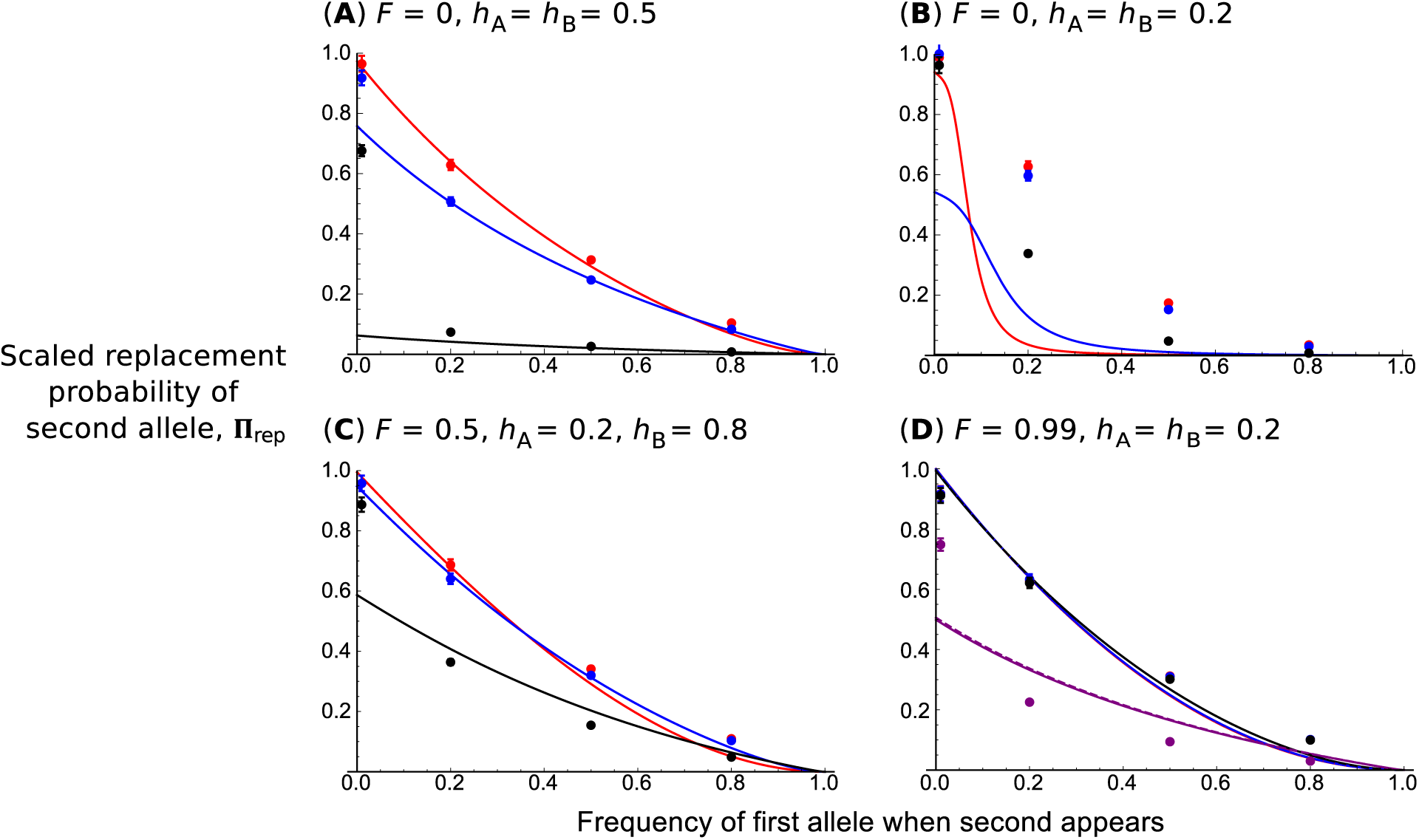
Probability Π_*rep*_ that a second beneficial allele with advantage *s_B_* replaces an existing sweep with selective advantage *s_A_*, where *s_B_* > *s_A_*, as a function of the first sweep frequency *p* when the second sweep appears. *N* = 5, 000 and *2Nr* = 0.01 (red), 0.1 (blue), 1 (black), or 100 (purple; panel (d) only). In (d), dashed lines are analytical results using the more exact recombination rate, *r*(1 – 2*F* + Φ). Other parameters are indicated above each panel. Points corresponds to 5,000 stochastic simulations for which the second beneficial allele has fixed. If confidence intervals cannot be seen, they lie within the plotted points.

To calculate a more accurate replacement probability in this case, it would be necessary to explicitly account for the frequency of the neutral haplotype (carrying no beneficial alleles). In addition, it would be desirable to explicitly track drift effects as recessive beneficial mutations are present at a low frequency. Unfortunately it will probably be unfeasible to produce tractable analytical solutions in either scenario. Hence in subsequent analyses when *ϕ >* 1, we will focus on codominant or dominant mutations (*h* ≥ 1/2).

### Codominant case

Under codominance (*h_A_* = *h_B_* = 1/2), selfing has no effect on the single-locus fixation probabilities. Here, we can therefore analyse the effect of selfing on recombination only. Moreover, for this specific case, results can be directly obtained by rescaling haploid models. Equations 11 and 12 become:

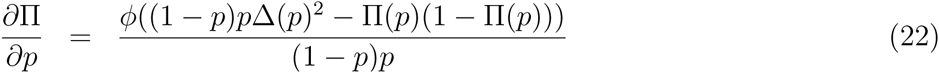

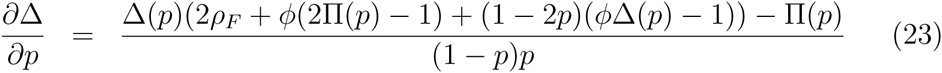

where *ρ*_*ρ*_ = *ρ*(1 – *F*)/(1 + *F*). Equations 22, 23 are similar to Barton’s (1995) 6a and 6b for haploids, except with (i) *p*(1 – *p*) terms in the denominator since our equations are as a function of the first sweep frequency, and (ii) that the recombination rate is decreased by 2(1 – *F*)/(1 + *F*). The latter scaling reflects how the population size is increased by a factor of 2 in diploids compared to haploids; how inbreeding magnifies drift by a factor 1/(1 + *F*), increasing the speed at which the first sweep fixes and reducing the potential for recombination to act; and how the effective recombination rate is reduced by 1 – *F* (Caballero and Hill 1992). Here, we can use the approximations given by Equations 8 and 9 a of Barton (1995) with the appropriate rescaling to find simplified forms for the total reduction in the relative fixation probability:

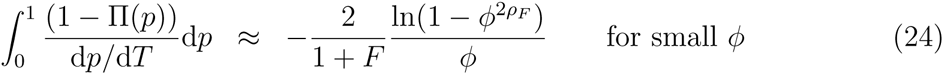

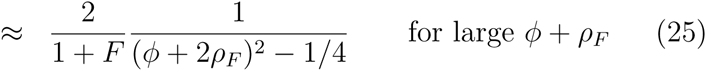

Approximations for the replacement probability can also be obtained (see details in Supplementary File S1). Quantitative inspection of previous equations shows that the emergence effect (or ‘stochastic interference’ effect) is more important than the replacement effect (Figure 4). The emergence effect is higher for low *ϕ* values, and can be very high; Equation 24 tends to infinity when *ϕ* or *ρ_F_* tend towards 0. On the contrary, *R̅* for the replacement case tends towards a small, finite value as *ϕ* tends towards infinity. This difference appears because (i) mutations are more sensitive to interference in the stochastic zone than once they have emerged, and (ii) selection interference is more pronounced when *ϕ* < 1 than when *ϕ* > 1 (and the second allele can replace the first). Consequently, the effect of selfing is more important for low *ϕ* values when emergence is the most important process, than for high *ϕ* values when replacement predominates, as illustrated in Figure 4. This figure also illustrates how the effect of a sweep can extend across long chromosome tracts with high selfing rates.

**Figure 4:**
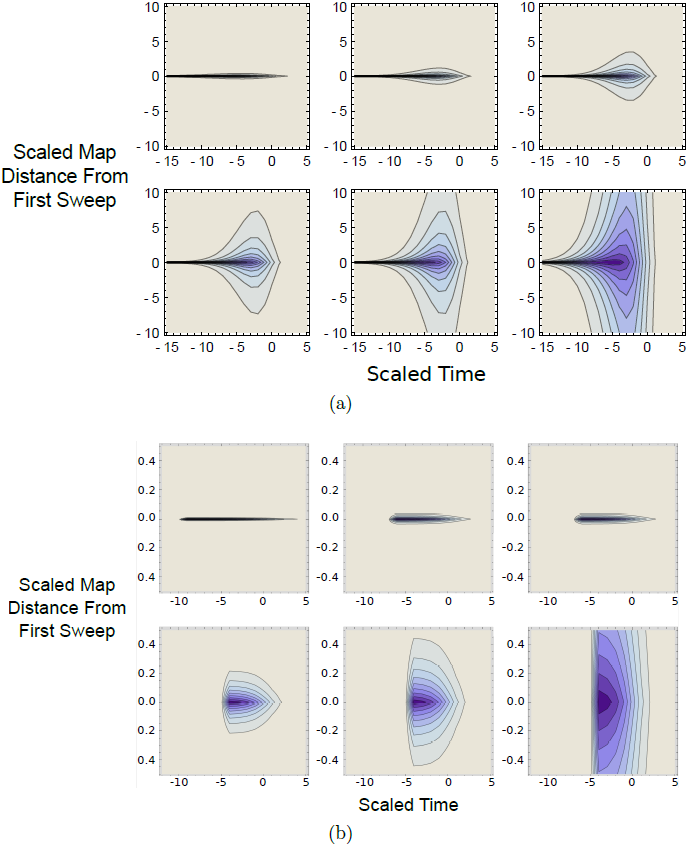
Contour plots showing degree of interference, as measured by Equation 2 with Π defined by Equation 22 (for *ϕ* < 1) and Π_*rep*_ defined with Equation 15 in Supplementary Material 1 (for *ϕ* > 1). Both beneficial mutations are additive (*h_A_* = *h_B_* = 1/2). In both panels, darker colours indicate higher degree of interference (with the darkest representing *R* approaching 0). x-axis denotes time of the sweep (with the sweep reaching 50% frequency at *T* = 0); y-axis is the map distance from the first sweep (scaled to 10^−2^/*s_A_*). Top row of plots are for *F* values of 0, 0.5, and 0.8 respectively; bottom row are *F* values of 0.9, 0.95, 0.99. Other parameters are *N* = 10, 000, and (a) *s_A_* = 0.01, *s_B_* = 0.005 so *ϕ* = 0.5; or (b) *s_A_* = 0.01, *s_B_* = 0.05 so *ϕ* = 5. Note difference y-axis scaling for (a) and (b).

In previous equations, the scaling factor 2/(1 + *F*) arises because the length of the sweep is in 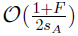, but we also scaled time by 1/*s_A_* to conserve the same scaling for any selfing rate. Equations 24 and 25 demonstrate the two opposite effects of selfing: the reduction in effective recombination reduces the probability of emergence, and also increases the replacement probability but over a shorter period of time as alleles fix more quickly (Glémin 2012). For loose linkage, the effect of selfing on recombination is stronger so that selfing globally decreases the probability of fixation. However, for tight linkage interference occurs for any selfing rate, such that the dominant effect of selfing is the reduction in sweep length (Figure A2 in Supplementary File S1). Boundary conditions can be found for the two extreme cases when *ϕ* is either small or large. When *ϕ* > 1, replacement is more likely under outcrossing than complete selfing if 4*Nr* < 1.386, (see Supplementary Mathematica File S2). When *ϕ* ≪ 1, emergence of the second beneficial allele is more likely under selfing than outcrossing only for very tight linkage, that is for *ρ* < –ϵ/4ln(*ϕ*), where ϵ is the residual outcrossing rate under selfing (see Supplementary Mathematica File S2).

### Effect of dominance on interference

For high selfing rates, the interference process is well approximated by the additive case. However, to get a complete picture of the effect of selfing we need to analyse how dominance affects the process. We will first consider outcrossing populations before turning to the effect of mating system on the adaptation rate. When investigating dominance, two questions arise. Which kind of mutations cause the strongest interference, and which ones are the most sensitive to interference and are hence more likely to be lost?

The effect of interference for different combinations of dominance levels are presented in Figure 5 for *ϕ* < 1. The main difference in sweep dynamics arises from the length of the two stochastic phases. Because a mutation causes interference mainly during its deterministic trajectory, which is similar for any dominance level (***O***(1/2*Ns_A_*) for any *h_A_*; Ewing *et al*. (2011)), the dominance level of mutation *A*_1_ has thus only a weak effect on the emergence probability of mutation *B*_1_. However, the sensitivity of mutation *B*_1_ to interference strongly depends on its dominance level, as it depends on the length of its initial stochastic phase, which is 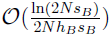 (Ewing *et al*. 2011). Recessive mutations are thus more sensitive to interference than additive and dominant ones. Interference thus reinforces Haldane’s Sieve, in the sense that recessive mutations are even less likely to emerge in outcrossing populations if tightly linked to the initial mutation.

**Figure 5:**
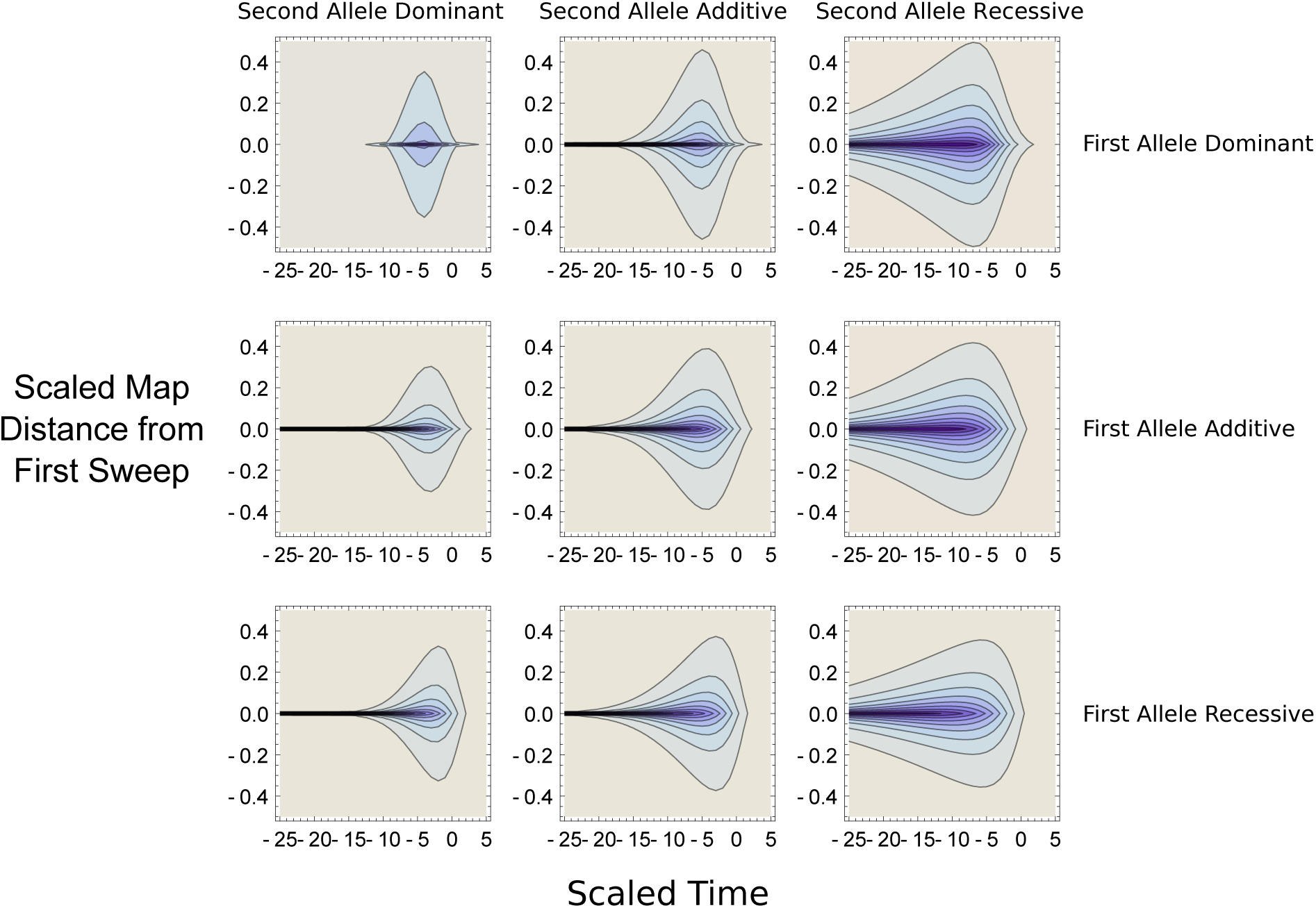
Contour plots showing degree of interference, as measured by Equation 2 with Π defined by Equation 22 and Π_*rep*_ = 0 (as *ϕ* < 1), for different dominance values. In both panels, darker colours indicate higher degree of interference (*R* approaching 0); x-axis denotes time of the sweep (with the sweep reaching 50% frequency at *T* = 0); y-axis is the map distance from the first sweep (scaled to 10^−2^/*s_A_*). Labels denote the dominance value of the first and second mutation, with recessive mutants having *h* = 0.2; additive mutations *h* = 0.5; dominant mutations *h* = 0.8. Other parameters are *N* = 10, 000, and *s_A_* = 0.01, *s_B_* = 0.005 so *ϕ* = 0.5.

In the case of strong interference, this effect can be substantial as illustrated in Figure 5. Interestingly, this effect is not symmetrical since dominant mutations only exhibit slightly less interference than additive mutations. As far as we know this effect has not been described before and leads to the prediction that the dominance spectrum of fixed beneficial mutations (i.e. the expected density of dominance values observed in fixed alleles) should vary with recombination rates (Figure A3 in Supplementary File S1).

### Conditions under which selection is more efficient under outcrossing than under selfing

We now have all the ingredients to study the range of conditions under which selfing reduces the adaptation rate. Without interference and other factors increasing drift in selfers, selfing reduces (respectively increases) adaptation from new dominant (respectively recessive) mutations (Charlesworth 1992; Caballero and Hill 1992). How does interference affect this behaviour? This question can be explored by considering a steady flow of mutations and analysing *P_AB_* = *R̅P_B_* where *R* is given by Equation 16. As shown in Supplementary Mathematica File S2, the total effect of interference on replacement will be no more than of the order of ln(2*Ns_A_*) (which is always lower than few tens) while the effect on emergence can be much more important. In what follows we will therefore focus on the case where *ϕ* < 1.

Figure 6 illustrates how selfing can affect the probability of fixation of the second mutation compared to the single locus case. Under a low adaptation regime (4*Nu_A_* = Θ = 0.02, for *u_A_* the per-locus beneficial mutation rate) interference is weak and the probability of fixation is reduced only in highly selfing species. This reduction is moderate and selfing species are still better than outcrossers at fixing recessive mutations. Under a stronger adaptation regime (Θ = 0.2), interference can be substantial even in mixed mating species and adaptation can be fully impeded in highly selfing species if 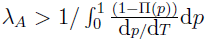 (see Barton (1995)). This threshold depends on *ϕ*, which means that, even under a low adaptation regime, weakly beneficial mutations can be affected by interference in highly selfing species. Figure 7 shows the joint dominance and selection spectrum for which selection is more efficient in outcrossing than in highly selfing (*F* = 0.95) species. Strongly beneficial mutations are very weakly affected by interference, so only dominant mutations are more efficiently selected in outcrossing than in selfing species. However, (very) weak beneficial mutations are better fixed in outcrossing populations, whatever their dominance level.

**Figure 6:**
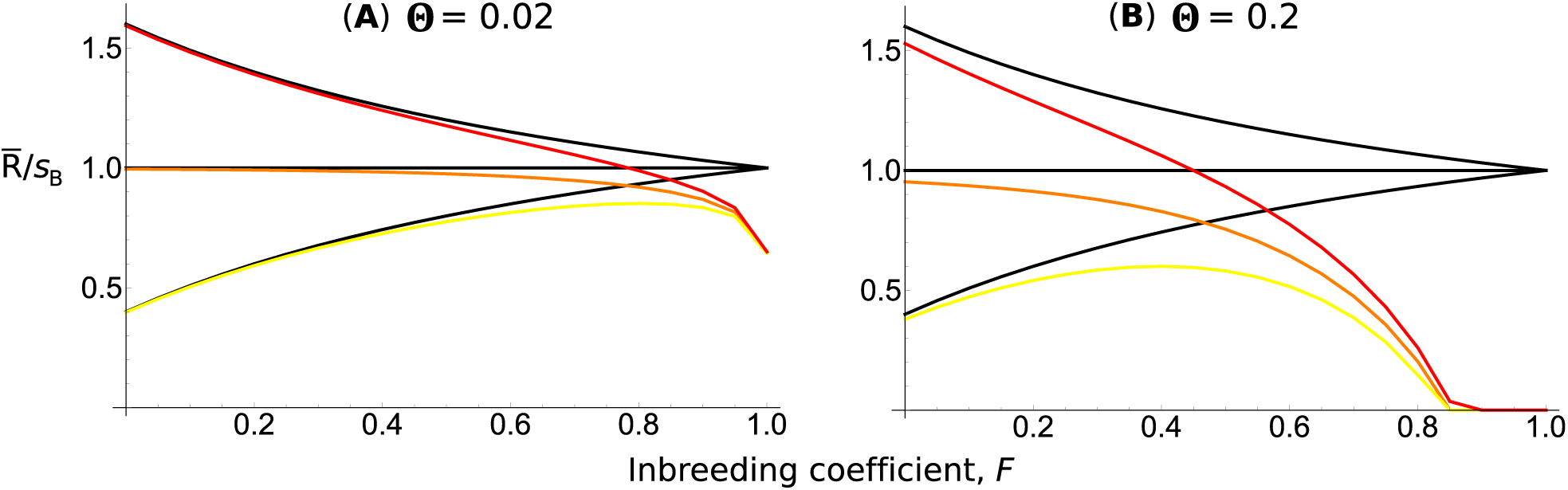
Plots of the total effect of interference, *R̅*, as defined using Equation 16, as a function of *F*. The *y*-axis is the probability of emergence scaled to *s_B_*, the expected emergence probability with *F* = 1. There is a continual rate of mutation Θ = 4*Nu_A_* = 0.02 (left) or 0.2 (right). *N* = 10, 000, *r* = 0.01, *h_A_* = 0.5, *s_A_* = 0.01, *s_B_* = 0.001 (ϕ = 0.1), and *h_B_* = 0.2 (yellow line), 0.5 (orange line), or 0.8 (red). Black lines show expected fixation probability in the absence of interference.

**Figure 7:**
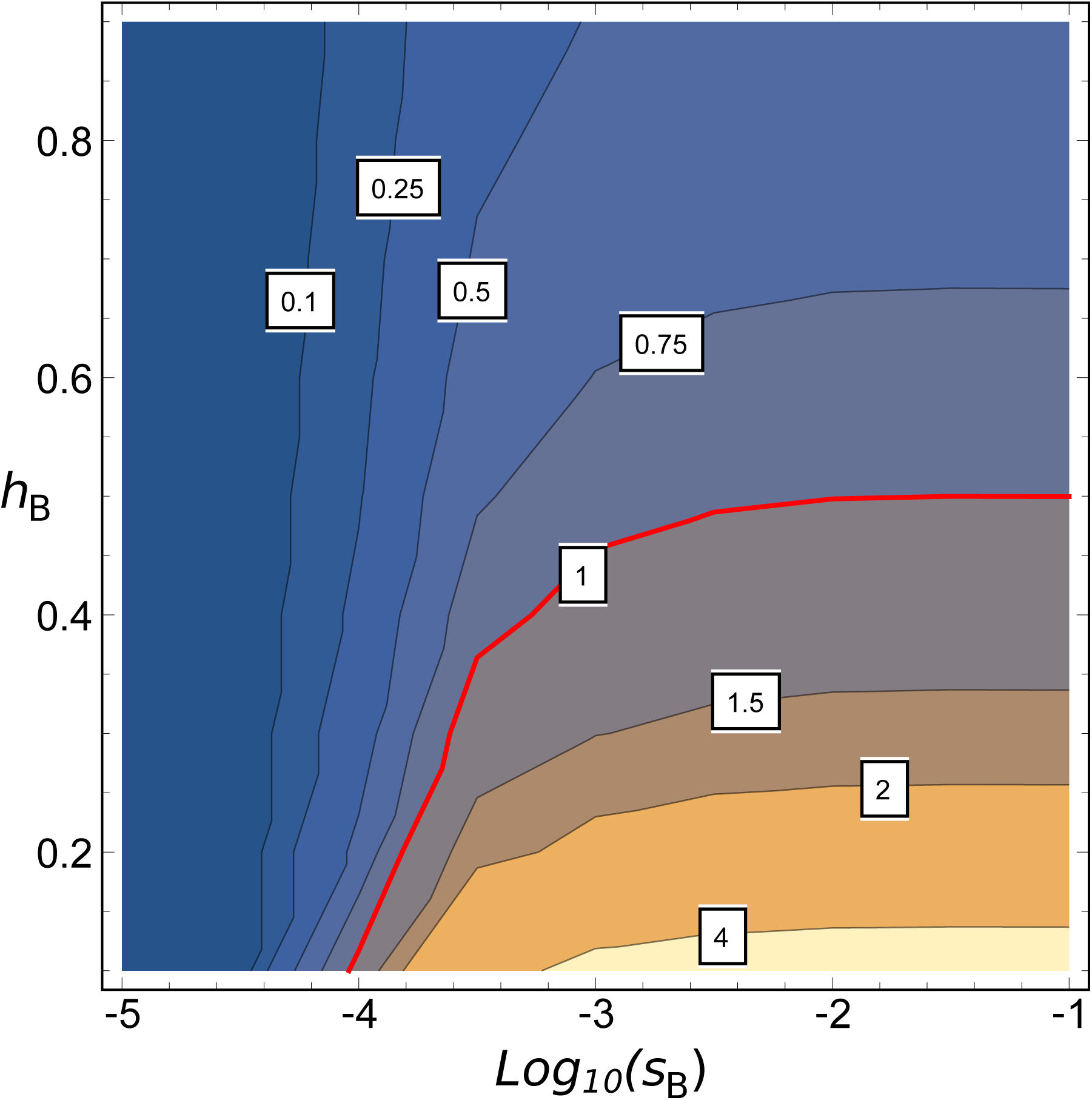
Contour plot of the ratio of *R̅* (Equation 16) for *F* = 0 and *F* = 0.95, as a function of *s_B_* (on a *log*_10_ scale) and *h_B_*. Values less than one indicate that outcrossers has the higher fixation probability, and values greater than one indicate that *F* = 0.95 populations have the higher probability. Other parameters are Θ = 0.1, *N* = 10, 000, *r* = 0.01, *h_A_* = 0.5, and *s_A_* = 0.01.

## Discussion

### Interference between beneficial mutations with partial selfing and dominance

Multi-locus models of adaptation in partial self-fertilising species can inform on how the interplay between homozygote creation, and reduction in recombination, jointly affect selection acting on multiple sites. It is already known that the presence of linked deleterious variation means that mildly recessive beneficial mutations (*h* just less than 1/2) are more able to fix in outcrossers than in selfing organisms by recombining away from the deleterious allele, in contrast to single locus theory (Hartfield and Glémin 2014). More generally, genome wide background selection can substantially reduce adaptation in highly selfing species (Kamran-Disfani and Agrawal 2014). Yet the extent that linkage between beneficial mutations impacts upon mating-system evolution remains poorly known.

Here we extended several previous models of selection interference to consider how adaptation is impeded in partially-selfing organisms. We considered two possibilities. First, given that an existing sweep is progressing through the population, subsequent mutations confer a lesser selective advantage and can only fix if recombining onto the fitter genetic background (the ‘emergence’ effect). Alternatively, a second mutant could be fitter and replace the existing sweep, unless recombination unites the two alleles (the ‘replacement’ effect). We found that the emergence effect is generally stronger than the replacement effect, and is more likely to lead to loss of beneficial mutations (Figure 4).

Furthermore, selection interference has two opposite effects on Haldane’s Sieve. In mainly outcrossing populations (where it operates), Haldane’s Sieve is reinforced because recessive mutations are even more likely to be lost when rare compared to dominant ones, compared to single locus results. However, when comparing different mating systems, interference reduces or nullifies the advantage of selfing of not being affected by Haldane’s Sieve. Consequently, weakly-beneficial mutations are more likely to be fixed in outcrossers, irrespective of their dominance level (Figure 7). These findings thus contribute to a body of literature as to when the predictions of Haldane’s Sieve should break down, or otherwise be altered. Other examples include the fixation probability of mutations being independent from dominance if arising from previously deleterious variation (Orr and Betancourt 2001); more generally, outcrossers are more able to fix mutations with any dominance level compared to selfers if arising from standing variation, and when multiple linked deleterious variants are present (Glémin and Ronfort 2013). Conversely, dominant mutations can be lost in metapopulations due to strong drift effects (Pannell *et al*. 2005).

In our model we assumed that no more than two beneficial mutations simultaneously interfere in the population. However, even if mutation does not occur frequently enough to lead to multiple mutations interfering under outcrossing, the presence of a few sweeping mutations throughout a genome can jointly interfere in highly selfing species. Obtaining a general model of multiple substitutions in a diploid partially selfing population is a difficult task, but it is likely that the rate of adaptation would be further reduced compared to the two-locus predictions (as found in haploid populations by Weissman and Barton (2012)).

It is also of interest to ask whether our calculations hold with different types of inbreeding (such as sib mating). For a single unlinked mutant, Caballero and Hill (1992) showed how various inbreeding regimes determine the value of *F* used in calculating fixation probabilities (Equation 1). However, it is unclear how effective recombination rates will be affected. For example, Nordborg’s (2000) rescaling argument relies on the proportion of recombination events that are instantly ‘repaired’ by direct self-fertilisation; these dynamics would surely be different under alternative inbreeding scenarios. Further work would be necessary to determine how other types of inbreeding affect net recombination rates, and thus the ability for selection interference to be broken down.

### Causes of limits to adaptation in selfing species

We have already shown in a previous paper how adaptation can be impeded in low-recombining selfing species due to the hitch-hiking of linked deleterious mutations (Hartfield and Glémin 2014), with Kamran-Disfani and Agrawal (2014) demonstrating that background selection can also greatly limit adaptation. Hence the question arises as to whether deleterious mutations or multiple sweeps are more likely to impede overall adaptation rates in selfing species.

Background selection causes a general reduction in variation across the genome by reducing *N_e_* (Nordborg *et al*. 1996); here the overall reduction in emergence probability is proportional to *N_e_*/*N*, where *N_e_* is mediated by the strength and rate of deleterious mutations (Barton 1995; Johnson and Barton 2002), and thus affects all mutations in the same way. Because of background selection, selfing is thus expected to globally reduce adaptation without affecting the spectrum of fixed mutations. Similarly, adaptation from standing variation, which depends on polymorphism level, is expected to be affected by the same proportion (Glémin and Ronfort 2013). Alternatively, interference between beneficial mutations is mediated by *ϕ*, the ratio of the selection coefficients of the sweeps. For a given selective effect at locus *A*, weak mutations at locus *B* are thus more affected by interference than stronger ones, and the net effect of interference cannot be summarised by a single change in *N_e_* (Barton 1995; Weissman and Barton 2012). Because of selective interference, selfing is also expected to shift the spectrum of fixed mutations towards those of strong effects. Interestingly, Weissman and Barton (2012) showed that neutral polymorphism can be significantly reduced by multiple sweeps, even if they do not interfere among themselves. This suggests that in selfing species, adaptation from standing variation should be more limited than predicted by single-locus theory (Glémin and Ronfort 2013). Selective interference could thus affect both the number and type of adaptations observed in selfing species.

Reflecting on this logic, both processes should interact and we therefore predict that background selection will have a diminishing-returns effect. As background selection lowers *N_e_* then the substitution rate of beneficial mutations will be reduced (since it is proportional to *N_e_μ* for *μ* the per-site mutation rate), hence interference between beneficial mutations will subsequently be alleviated. No such respite will be available with a higher adaptive mutation rate; on the contrary, interference will increase (Figure 6). Impediment of adaptive alleles should play a strong role in reducing the fitness of selfing species, causing them to be an evolutionary dead-end. Further theoretical work teasing apart these effects would be desirable. Given the complexity of such analyses, simulation studies similar to those of Kamran-Disfani and Agrawal (2014) would be a useful approach to answering this question.

In a recent study, Lande and Porcher (2015) demonstrated that once the selfing rate became critically high, selfing organisms then purged a large amount of quantitative trait variation, limiting their ability to respond to selection in a changing environment. This mechanism provides an alternative basis as to how selfing organisms are an evolutionary dead-end. However, they only consider populations at equilibrium; our results suggest that directional selection should further reduce quantitative genetic variation due to selective interference among mutations. Subsequent theoretical work is needed to determine the impact of interference via sweeps on the loss of quantitative variation. Furthermore, complex organisms (i.e. those where many loci underlie phenotypic selection) are less likely to adapt to a moving optimum compared to when only a few traits are under selection (Matuszewski *et al*. 2014), and can also purge genetic variance for lower selfing rates (Lande and Porcher 2015). Complex selfing organisms should thus be less able to adapt to environmental changes.

### Empirical Implications

The models derived here lead to several testable predictions for the rate of adaptation between selfing and outcrossing sister-species. These include an overall reduction in the adaptive substitution rate in selfing populations; a shift in the distribution of fitness effects in selfing organisms to only include strongly-selected mutations that escape interference; and a difference in the dominance spectrum of adaptive mutations in outcrossers compared to selfers, as already predicted by single-locus theory (Charlesworth 1992) and observed with quantitative trait loci (QTLs) for domesticated crops (Ronfort and Glémin 2013).

So far, few studies currently exist that directly compare adaptation rates and potential between related selfing and outcrossing species, but they are in agreement with the predictions of the model. In plants, the self-incompatible *Capsella grandiflora* exhibited much higher adaptation rates (where *α* = 40% of non-synonymous substations were estimated to be driven by positive selection using the McDonald-Kreitman statistic; Slotte *et al*. (2010)) than in the related selfing species *Ara-bidopsis thaliana* (where *α* is not significantly different from zero). Similarly, the outcrossing snail *Physa acuta* exhibited significant adaptation rates (*α* = 0.54), while no evidence for adaptation in the selfing snail was obtained (Burgarella *et al*. 2015); in fact, evidence suggests that deleterious mutations segregate due to drift (*α* = –0.19). In agreement with the predicted inefficacy of selection on weak mutations, Qiu *et al*. (2011) also observed significantly lower selection on codon usage in the *Capsella* and *Arabidopsis* selfers than in their outcrossing sister species.

In addition, as only strong advantageous mutations are expected to escape loss through selection interference, this result can explain why selective sweeps covering large tracts of a genome are commonly observed, as with *Arabidopsis thaliana* (Long *et al*. 2013) and *Caenorhabditis elegans* (Andersen *et al*. 2012). Extended sweep signatures can also be explained by reduced effective recombination rates in selfing genomes. Finally, selective interference between beneficial mutations could explain why maladaptive QTLs are observed as underlying fitness components, as detected in *Arabidopsis thaliana* (Ågren *et al*. 2013). Direct QTL comparisons between selfing and outcrossing sister species would therefore be desirable to determine to what extent selection interference leads to maladaptation in selfing species.

## Acknowedgements

M.H. was funded by an ATIP-Avenir grant from CNRS and INSERM to Samuel Alizon, a Marie Curie International Outgoing Fellowship grant number MC-IOF-622936 (project SEXSEL), and also acknowledges additional support from the CNRS and the IRD. S.G. is supported by the French CNRS and a Marie Curie Intra-European Fellowship, grant number IEF-623486 (project SELFADAPT). This work was also supported by two grants from the Agence Nationale de la Recherche (TRANS: ANR-11-BSV7-013-03 and SEAD: ANR-13-ADAP-0011). We thank Joachim Hermisson, Denis Roze and an anonymous reviewer for their very helpful comments on the manuscript.

